# Cyclic-Phe-Pro Binds the ToxRS Interface to Promote Signal Transduction in *Vibrio vulnificus*

**DOI:** 10.64898/2026.07.20.739674

**Authors:** Tae-Yeon Kim, Woo-Chan Ahn, Jeong-A Kim, Keun-Woo Lee, Soyee Kim, Kwang-Hyun Park, Eui-Jeon Woo, Kun-Soo Kim

## Abstract

In *Vibrio vulnificus*, the quorum-sensing signal cyclo-(L-phenylalanine-L-proline) (cFP) binds the membrane receptor ToxRS to activate genes linked to oxidative-stress resistance and virulence. ToxR is a transmembrane transcription factor that pairs with ToxS to sense periplasmic signals, yet how *V. vulnificus* ToxRS recognizes cFP has remained undefined. AI-guided structure prediction revealed preferential ToxR/S heterodimer formation and a fold conserved with the *V. cholerae* crystal structure. Molecular docking placed cFP in a hydrophobic pocket at the ToxR–ToxS interface, where Phe279 stacked with its phenyl ring and Arg277 hydrogen-bonded its carbonyl oxygen. Introducing R277L and F279A substitutions lowered basal *leuO* expression and abolished cFP-dependent induction in a *lacZ* fusion assay. ChIP showed that cFP enhanced wild-type ToxR binding to the *leuO* promoter, whereas the mutant bound weakly and did not respond. Thus cFP bridges ToxR and ToxS to stabilize the heterodimer, facilitating ToxR recruitment to promoters and transcriptional activation in *V. vulnificus*.

## Introduction

*Vibrio vulnificus* is a halophilic gram-negative bacterium and an opportunistic pathogen that causes primary septicemia with a high mortality rate (Baker-Austin et al. 2018). To coordinate survival and virulence, *V. vulnificus* employs a quorum-sensing (QS) pathway mediated by the cyclic dipeptide cyclo-(L-phenylalanine-L-proline) (cFP) (Park et al. 2006; Kim et al. 2013a; Kim et al. 2013b; Kim et al. 2018; Park et al. 2019). Cells produce and secrete cFP, which re-enters neighboring cells by passive diffusion (Park et al. 2020). The inner membrane-anchored receptor complex ToxRS detects this signal and directly upregulates the expression of *ompU* and *leuO* (Park et al. 2006; Park et al. 2019). The *ompU*-encoded porin mediates host cell adhesion (Goo et al. 2006), while LeuO activates a downstream cascade involving the sigma factor RpoS and its target *katG*, whose catalase product detoxifies reactive oxygen species within the host (Kim et al. 2018). This cFP–ToxRS pathway collectively regulates more than 900 genes in *V. vulnificus* (Kim et al. 2013a).

cFP initiates the signaling by binding to the periplasmic domain of ToxRS (Park et al. 2019). ToxR is a transmembrane transcription factor comprising an N-terminal DNA-binding domain, a transmembrane segment, and a C-terminal periplasmic domain (ToxRp) (Miller et al. 1987). ToxR is co-expressed with ToxS, whose periplasmic domain (ToxSp) interacts with ToxRp to form a heterodimeric complex that stabilizes ToxR and enhances its transcriptional activity (DiRita and Mekalanos 1991). The oligomeric state of ToxR has been debated; early studies proposed a homodimer model, but this form depends on cysteine oxidation under non-physiological over-expression conditions (Ottemann and Mekalanos 1996). The crystal structure of the *V. cholerae* ToxRpSp complex confirmed the heterodimer as the physiological signaling unit (Gubensak et al. 2023). Importantly, *V. cholerae* ToxRS senses bile acid (Martinez-Hackert and Stock 1997; Midgett et al. 2017), whereas *V. vulnificus* ToxRS responds to cFP, a cyclic dipeptide produced as a quorum sensing signal (Park et al. 2006; Park et al. 2019). cFP accumulates in a cell-density-dependent manner and activates a downstream cascade ToxR → LeuO → vHUαβ → RpoS → KatG promoting virulence (Kim et al. 2018). ToxRp and ToxSp of *V. vulnificus* share only 55% and 71.5% amino acid identity with their *V. cholerae* counterparts (Lee et al. 2000). The structural basis of cFP recognition by *V. vulnificus* ToxRS therefore remains unknown.

Deep learning-based structure prediction tools have substantially advanced the ability to model protein complexes that are difficult to resolve experimentally (Jumper et al. 2021; Abramson et al. 2024). The difficulty in obtaining a ToxRS structure arises not from membrane anchoring alone, but from the conformational instability of the ToxR periplasmic domain (ToxRp). The C-terminus of ToxRp is largely disordered in isolation and folds into ordered structure only upon binding ToxS (Gubensak et al. 2021; Gubensak et al. 2023). Consistent with this, an intra-chain disulfide bond is required to constrain a flexible loop and stabilize the ToxS-binding region of ToxRp (Midgett et al. 2020). Prior computational analyses predicted ToxRpSp structures from diverse *Vibrio* species and confirmed that the overall heterodimeric fold is well conserved, with the ToxSp β-barrel being especially invariant across species (Gubensak et al. 2023). The conserved residues within the ToxSp binding pocket were proposed to accommodate small hydrophobic ligands broadly similar to bile acids. However, no comparative docking analysis has been performed to identify cFP-specific binding determinants in *V. vulnificus* ToxRS, leaving the molecular basis of ligand discrimination between species unaddressed. Integrating structure prediction with molecular docking and site-directed mutagenesis offers a practical strategy for generating and testing mechanistic hypotheses in systems where experimental structural data are unavailable.

In this study, we used deep learning-based structural modeling to predict the three-dimensional structure of the *V. vulnificus* ToxRpSp complex and to compare the stability of ToxR/S heterodimer versus ToxR/R homodimer formation. Molecular docking identified a cFP-binding pocket at the ToxR–ToxS interface, with Arg277 (R277) and Phe279 (F279) of ToxR as the predicted key contact residues. We introduced R277L and F279A substitutions and evaluated their functional consequences using *leuO*-*lacZ* transcriptional fusion assays and chromatin immunoprecipitation (ChIP) analysis. Our results demonstrate that R277 and F279 are essential for cFP-mediated signaling *in vivo*, and support a model in which cFP binds at the ToxR–ToxS interface to enhance heterodimer formation and facilitate transcriptional activation of target genes in *V. vulnificus*.

## Materials and Methods

### Strains, culture conditions, and chemicals

The bacterial strains and plasmids used in this study are listed in Table S1. *Vibrio vulnificus* strains were cultured in Luria-Bertani (LB) broth with appropriate antibiotics or in thiosulfate citrate bile salt sucrose (TCBS) agar at 30°C. *Escherichia coli* strains were cultured in LB broth with appropriate antibiotics at 37°C. All primers are listed in Tables S1 and S2. cFP (Bachem Inc., Switzerland) was dissolved in DMSO and used at a final concentration of 5 mM.

### Ligand binding prediction to ToxRpSp heterodimer

The complex structure of ToxRp and ToxSp was modeled using AlphaFold3 (Abramson et al. 2024), and the most optimal structure was selected. Prior to docking, the protein and ligand structures were preprocessed using OpenBabel (O’Boyle et al. 2011). Molecular docking was performed using AutoDock Vina v1.2.3 (Trott and Olson 2010; Eberhardt et al. 2021) with a docking grid size of 15 Å × 15 Å × 16 Å. Structural visualization was carried out using PyMOL (Schrodinger 2015).

### Construction of pBBR1-MCS2::*toxRS*

All the primers used in this study are listed in Supplementary Table S2. A 1643-bp DNA fragment comprising the promoter region and the coding region of *toxRS* was amplified by PCR using primers toxRS_comp_F and toxRS_com_R and ligated into the *Bam*HI and *Eco*RI sites of pBBR1-MCS2 (Kovach et al. 1995) to generate pBBR12::*toxRS*.

### Site-directed mutagenesis of ToxRp

For site-directed mutagenesis of the cFP binding site, a 5,953-bp DNA fragment was amplified from pBBR12::*toxRS* by PCR using the primers, toxR_SDM_F and toxR_SDM_R, and PrimeSTAR^®^ HS DNA Polymerase (Takara, Tokyo, Japan). The amplified product was digested with restriction enzyme *Dpn*I (Enznomics, Daejeon, Korea) to remove original templates which were then transformed into *E. coli* S17-1 (Simon et al. 1983). The resulting construction, pBBR1-MCS2::*toxR*_*mt*_*S* contains R277L and F279A mutations.

*E. coli* S17-1 strains containing either pBBR1-MCS2::*toxRS* or pBBR1-MCS2::*toxR*_*mt*_*S* were mobilized into the MO6-24/O strain (Park et al. 2019) by conjugation to obtain the strain Δ*toxRS*.

### Construction of *leuO-lacZ* transcription fusions and β-galactosidase assay

A *lacZ* transcription fusion vector, pRK-*leuO*::*lacZ* (Park et al. 2019) was conjugated into appropriate derivatives of MO6-24/O using *E. coli* S17-1 and confirmed as *V. vulnificus* w ith TCBS agar.

β-Galactosidase activity from cells harboring the genes fused with *lacZ* vector described above was measured as described previously (Miller 1972). Briefly, derivatives of Δ*toxRS* cells were grown overnight in LB medium and sub-cultured into fresh LB medium at an initial OD_600_ of 0.005. To determinate the effect of cFP, each broth was supplemented with either 5mM of cFP or 12.5% DMSO as a negative control.

### Expression and purification of the periplasmic domains of ToxR and ToxR_mt_/S

C-terminal periplasmic domains of ToxRp (98 aa) and ToxSp (153 aa) were cloned into pCold− I (Takara, Tokyo, Japan) to yield pColdI::toxRp and pColdI::toxSp; pColdI::toxRmtp was constructed by site-directed mutagenesis. His-tagged proteins expressed in E. coli BL21(DE3) via IPTG cold-shock induction were purified by Ni-NTA chromatography (Qiagen) and checked by SDS-PAGE. Anti-ToxSp polyclonal antibody was prepared per (Kim et al. 2015).

### ChIP analysis

ChIP was performed per the manufacturer’s protocol (EZ ChIP, Upstate Biotechnology). Cells (OD_600_ ≈ 2.0) were crosslinked via 1% formaldehyde, quenched with 0.125 M glycine, and sonicated. After overnight immunoprecipitation at 4°C with anti-ToxR polyclonal antibody (AbClon, Seoul, Korea) or control IgG, purified DNA was quantified by qPCR (CFX Opus 96, Bio-Rad) using leuO_ChIP_F/R primers (Table S2).

## Results

### AlphaFold3 predicts preferential heterodimer formation of *V. vulnificus* ToxRS over ToxR homodimer

To understand the structural basis of cFP signal transduction, we predicted the three-dimensional structure of the *V. vulnificus* ToxRpSp complex using AlphaFold3 (Abramson et al. 2024). The predicted structure adopts the conserved ToxRS periplasmic architecture, in which ToxSp folds into a compact β-barrel and ToxRp packs against one face through a mixed α/β surface to form the heterodimer (Fig. 1a). The model showed high structural confidence, with a mean predicted local distance difference test (pLDDT) score of 92.02. The interchain predicted aligned error (pAE), which reflects confidence in the relative orientation of two chains at the interface, was 5.82 Å for the ToxRp/Sp heterodimer (Fig. S1), compared with 8.93 Å for the predicted ToxRp/Rp homodimer. The lower pAE for the heterodimer indicates that AlphaFold3 predicts a more confident and well-defined interchain arrangement, consistent with physiological observations in related Vibrio species. (Ottemann and Mekalanos 1996; Gubensak et al. 2021)

**Fig. 1.**
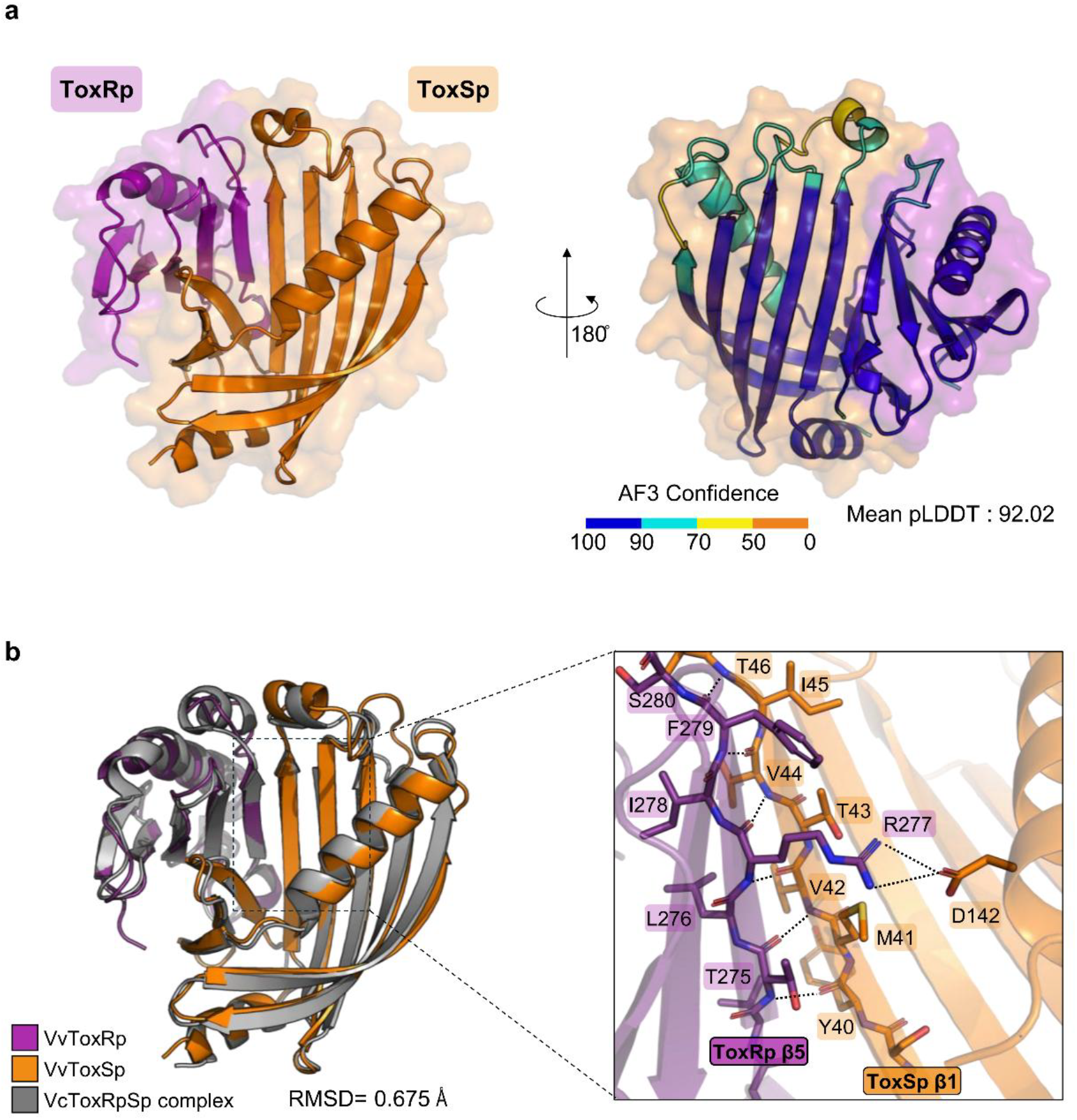
AlphaFold3 prediction of the *V. vulnificus* ToxRpSp heterodimer and comparison with the *V. cholerae* ToxRpSp complex. (a) Predicted structure of the periplasmic ToxR–ToxS heterodimer from *V. vulnificus*. ToxRp and ToxSp are shown in purple and orange, respectively, in two orientations rotated by 180°. The AlphaFold3 confidence score is indicated by the color scale, and the predicted complex showed a mean pLDDT score of 92.02. (b) Structural superposition with the *V. cholerae* ToxRpSp crystal structure (PDB: 8ALO; gray). The magnified view highlights the ToxRp β5–ToxSp β1 interface Residues Thr275, Leu276, Arg277, Ile278, Phe279, and Ser280 of ToxRp and Tyr40, Met41, Val42, Thr43, Val44, Ile45, Thr46, and Asp142 of ToxSp are indicated. Dashed lines denote predicted interchain contacts.

The predicted *V. vulnificus* ToxRpSp structure showed an RMSD of 0.675 Å against the *V. cholerae* crystal structure (PDB: 8ALO), confirming a highly conserved overall fold (Fig. 1b). At the heterodimer interface, ToxRp β5 (residues 274–278) forms a main-chain hydrogen bond network with ToxSp β1 (residues 41–45), completing the eight-stranded β-barrel of ToxSp. An intermolecular salt bridge between Arg277 of ToxRp and Asp142 of ToxSp further stabilizes the interface at the barrel opening.

### Molecular docking reveals key interactions of cFP with the ToxRS heterodimer interface

Molecular docking using AutoDock Vina identified a cFP binding pocket at the ToxRpSp heterodimer interface (Fig. 2a). The top-ranked pose yielded a binding affinity of −8.123 kcal/mol. The top four poses clustered within an RMSD of 2.0 Å from the best mode (RMSD l.b.: 1.41–1.95 Å), indicating convergence of the predicted binding geometry. All subsequent interaction analyses are based on the best-ranked pose.

**Fig. 2.**
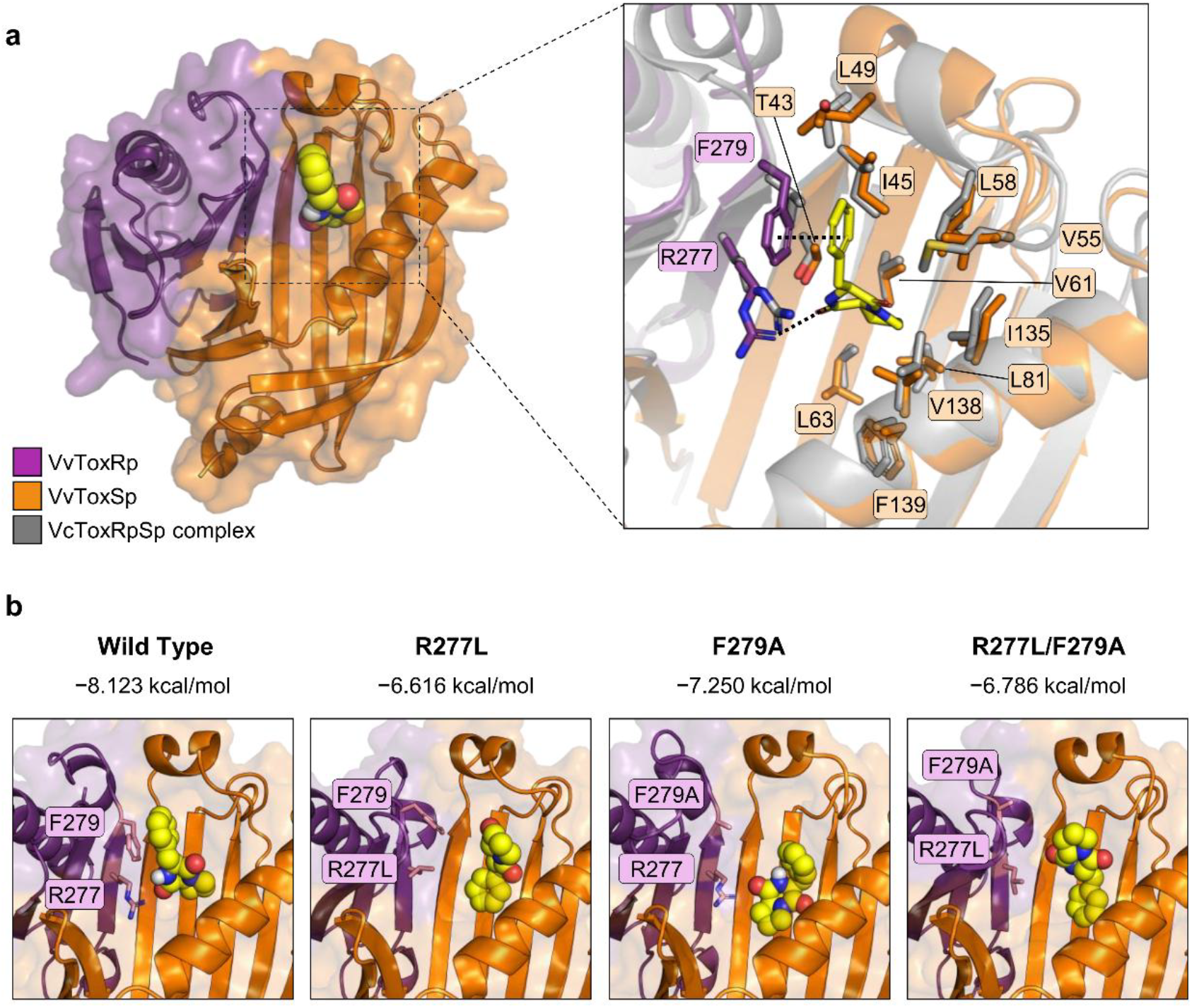
Predicted binding mode of cFP at the ToxRp–ToxSp heterodimer interface. (a) Molecular docking of cFP with the predicted ToxRp–ToxSp heterodimer. ToxRp and ToxSp are shown in purple and orange, respectively. cFP is shown as yellow spheres (overview) and yellow sticks (magnified view). Key contact residues Arg277 and Phe279 of ToxRp and hydrophobic pocket residues of ToxSp are labeled. The corresponding *V. cholerae* ToxRpSp structure (PDB: 8ALO; gray) is superposed for comparison. Dashed lines denote predicted interactions. (b) Docked poses of cFP (yellow spheres) at wild-type and mutant (R277L, F279A, and R277L/F279A) interfaces. Top-ranked binding affinities from AutoDock Vina are indicated above each panel. Mutated residues are shown in pink.

The binding pocket is predominantly hydrophobic, lined by ToxSp residues Ile45, Leu49, Val55, Leu58, Val61, Leu63, Leu81, Leu83, and Ile94, the majority of which are conserved as hydrophobic residues across *Vibrio* species. Within this cavity, the phenyl ring of cFP engages in face-to-face π–π stacking with Phe279 of ToxRp at a distance of 3.6 Å. Additionally, the guanidinium NH of Arg277 of ToxRp forms a hydrogen bond with the carbonyl oxygen of cFP at 2.8 Å. The involvement of both ToxRp residues (Arg277, Phe279) and ToxSp residues in coordinating cFP confirms that the ligand occupies the interface between the two proteins, consistent with a role for cFP in stabilizing the heterodimeric complex.

To examine the functional roles of Arg277 and Phe279, we introduced two substitutions: R277L, which eliminates the guanidinium-mediated hydrogen bond, and F279A, which removes the aromatic side chain responsible for π–π stacking. AlphaFold3 modeling confirmed that neither mutation impairs heterodimer formation (ipTM 0.85 for R277L/F279A vs. 0.83 for wild type). AutoDock Vina docking showed reduced cFP binding affinity for all mutants compared to wild type (−8.123 kcal/mol): −7.250 kcal/mol for F279A, −6.786 kcal/mol for R277L/F279A, and −6.616 kcal/mol for R277L (Fig. 2b), with more dispersed docking poses reflecting a less defined binding geometry.

### The *t*oxRmt mutant showed decreased expression of *leuO*

To verify the functional importance of the predicted binding site, we generated a mutant strain, ToxRmt, containing R277L and F279A substitutions. We then measured the transcription rate of the target gene, *leuO*, using a *lacZ* fusion assay. In wild-type cells (pBBR12::*toxRS*), the addition of 5 mM cFP significantly increased *leuO* expression, reaching approximately 12,000 Miller Units. In contrast, the *ToxR*_*mt*_ strain showed a significantly lower basal level of *leuO* expression (approx. 5,000 Miller Units), and the induction effect of cFP was almost entirely abolished (Fig. 3). The results demonstrate that the R277 and F279 residues are essential for cFP-mediated signal transduction.

**Fig. 3.**
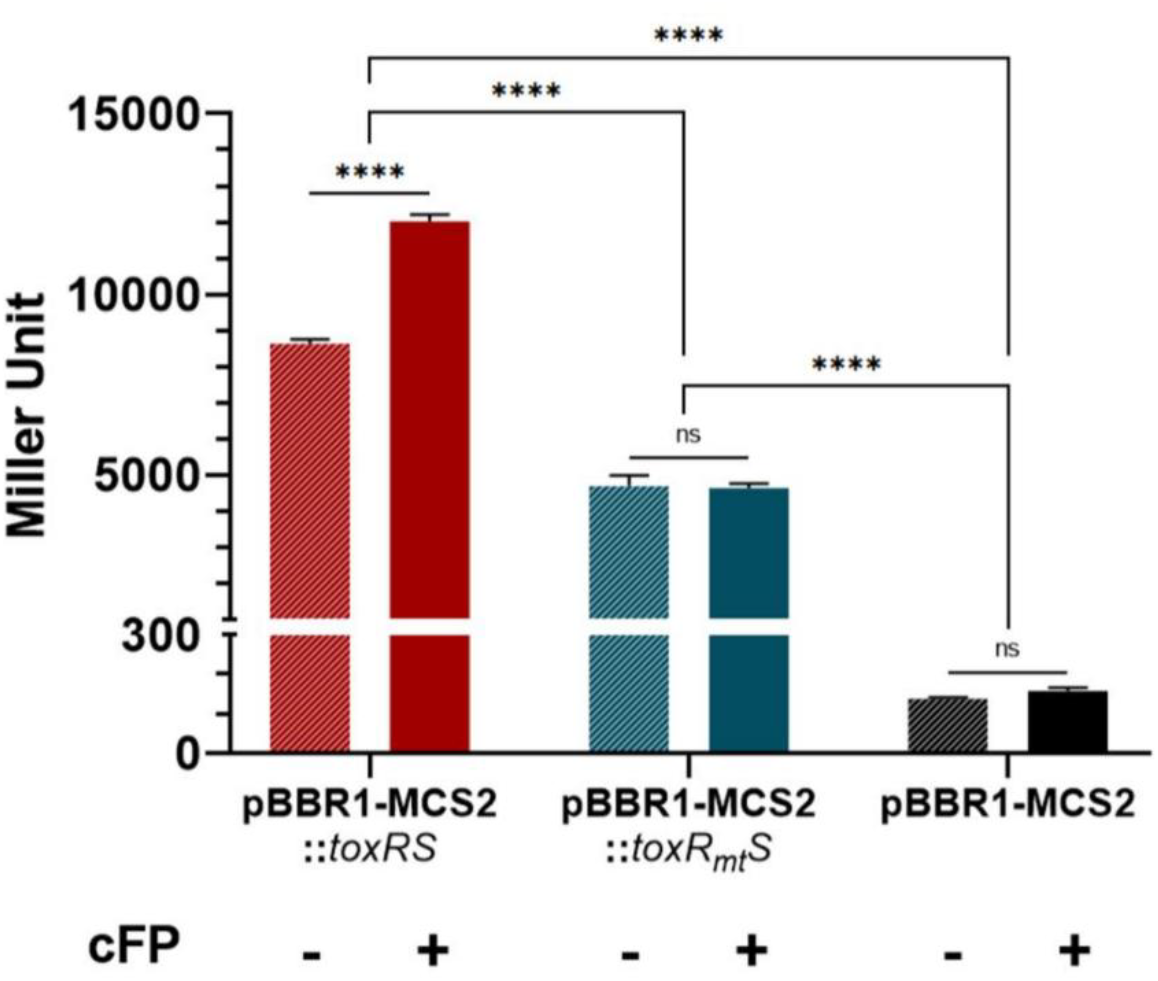
Effect of ToxR R277L/F279A mutations on the transcription of *leuO* as measured using the *lacZ*-transcriptional fusion to *leuO*. β-Galactosidase activity of the P*leuO-lacZ* transcriptional fusion was measured in the *V. vulnificus* ΔtoxRS strain harboring pBBR1-MCS2::*toxRS*, pBBR1-MCS2::*toxRmtS*, or the empty vector pBBR1-MCS2. The *toxRmtS* construct encodes ToxR carrying R277L and F279A substitutions. Cells were grown in the absence or presence of 5 mM cFP. Promoter activity is expressed as Miller Units. Data are shown as the mean ± SD. Statistical significance is indicated above the bars. ****, P < 0.0001; ns, not significant.

### cFP facilitates the binding of ToxR onto the upstream region of *leuO* but *ToxR*_*mt*_ does not

Finally, we conducted Chromatin Immunoprecipitation (ChIP) assays to confirm whether cFP promotes the recruitment of *ToxR* to the *leuO* promoter *in vivo*. In the presence of cFP, wild-type *ToxR* showed a 6-fold to 32-fold increase in enrichment at the *leuO* promoter region. The ToxR_*mt*_/S complex exhibited significantly lower affinity for the promoter compared to the wild-type, and its binding was not enhanced by cFP (Fig. 4). This confirms that cFP-induced stabilization of the ToxR/S heterodimer is a prerequisite for effective DNA binding and subsequent transcriptional activation.

**Fig. 4.**
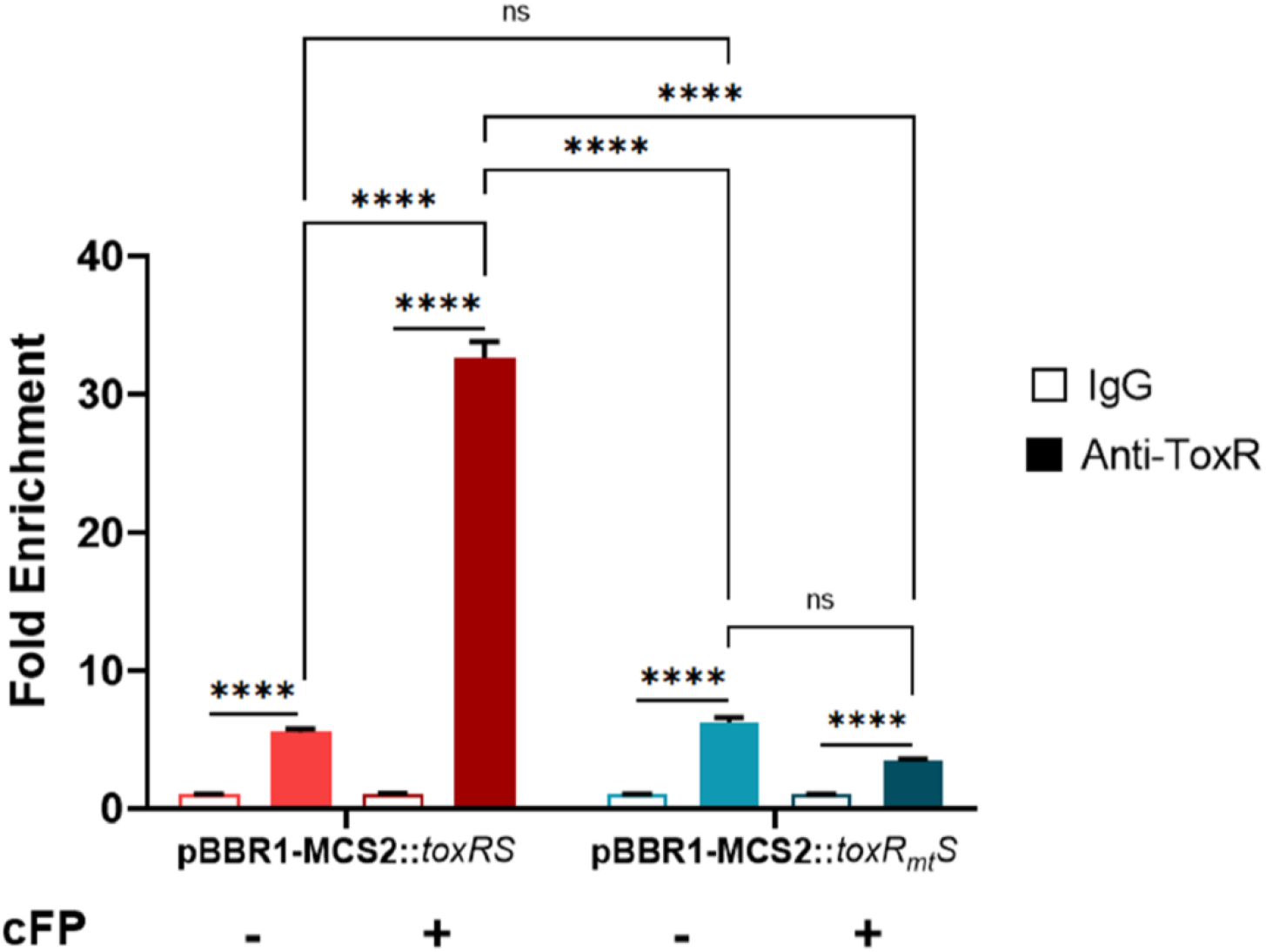
ChIP-qPCR analysis of ToxR binding to the *leuO* promoter region. Chromatin immunoprecipitation followed by quantitative PCR was performed using anti-ToxR antibody or control IgG. The *V. vulnificus* ΔtoxRS strain harboring pBBR1-MCS2::*toxRS* or pBBR1-MCS2::*toxRmtS* was grown in the absence or presence of 5 mM cFP. Enrichment of the *leuO* promoter region is shown as fold enrichment. Data are shown as the mean ± SD. Statistical significance is indicated above the bars. ****, P < 0.0001; ns, not significant.

## Discussion

### Structural insight into the ToxR/S heterodimerization

The oligomeric state of ToxR has been debated, with the homodimer now regarded as a non-physiological artifact of cysteine oxidation under overexpression (Ottemann and Mekalanos 1996; Gubensak et al. 2021). Our AlphaFold3 modeling supports the heterodimer as the favored assembly: the ToxRp/Sp interface was predicted with high confidence (pAE 5.82 Å), whereas the ToxRp/Rp homodimer was not (pAE 8.93 Å) (Fig. S1). The close fit to the *V. cholerae* structure (RMSD 0.675 Å) indicates a conserved fold (Fig. 1b), justifying cross-species comparison while leaving room for species-specific ligand recognition. This implies that signaling in *V. vulnificus* is governed not by ToxR concentration alone but by its specific interaction with ToxS, positioning the heterodimer as the unit that senses environmental cues.

### In silico prediction of the cFP binding mechanism

Molecular docking with AutoDock Vina placed cFP in a hydrophobic pocket at the ToxRp– ToxSp interface rather than within either protomer (−8.123 kcal/mol), with converged poses indicating a defined geometry (Fig. 2a). The pocket is lined by conserved ToxSp residues, while two ToxRp residues make the specific contacts: Phe279 stacks with the cFP phenyl ring (3.6 Å) and Arg277 hydrogen-bonds its carbonyl oxygen (2.8 Å). Because the ligand engages both protomers, this model casts cFP as a “molecular glue” that bridges ToxR and ToxS and thereby promotes their dimerization. Modeling the R277L and F279A substitutions reinforced this assignment. The mutant complexes retained interface confidence comparable to the wild type (ipTM 0.85 vs 0.83), yet the predicted cFP affinity fell to between −6.616 and −7.250 kcal/mol and the docked poses no longer converged (Fig. 2b). This computational separation of effects indicates that Arg277 and Phe279 act as determinants of cFP recognition rather than of heterodimer assembly.

### cFP stabilizes the complex to drive transcription

Our findings couple cFP perception directly to stabilization of the heterodimer. Because the substitutions leave the predicted heterodimer intact, the in vivo defects can be attributed to loss of cFP recognition rather than to a collapse of the complex. In the absence of cFP, the complex may lack the rigidity or orientation required for optimal activity; cFP binding enhances its formation and stability and, in turn, the recruitment of ToxR to the *leuO* promoter. The mutational data support this: R277L/F279A lowered basal *leuO* transcription and abolished cFP induction (Fig. 3), and reduced ToxR occupancy at the promoter while eliminating its cFP-dependent increase (Fig. 4). The increased DNA-binding seen by ChIP indicates that this structural stabilization is a prerequisite for transcriptional activation, and the loss of responsiveness precisely when the predicted contacts are removed links the docking prediction to the in vivo phenotype.

### Biological implications

This mechanism lets *V. vulnificus* convert a cyclic dipeptide signal into a decisive transcriptional switch. Coupling signal sensing to heterodimer stability provides a ligand-gated, switch-like response. The cFP–ToxRS axis acts at the top of the LeuO regulatory pathway and shapes the expression of hundreds of genes (Kim et al. 2013a; Kim et al. 2018). This ligand-dependent control thus links population density to oxidative-stress resistance and virulence.

## Statements and Declarations

## Acknowledgements

This work was supported by National Research Foundation of Korea (NRF) grants funded by the Korean Government (MSIP) (NRF-2019R1A2C2084282 to KKS; and RS-2022-NR071772, RS-2021-NR059435 and RS-2021-NR056526 to WEJ), the National Research Council of Science & Technology (NST) (CRC22024-500 to WEJ), and the KRIBB Research Initiative Program (KGM5382632, KGM1062612, KGM1322612 to WEJ).

## CRediT authorship contribution statement

Tae-Yeon Kim: Investigation, Formal analysis, Writing – original draft, Writing – review & editing. Woo-Chan Ahn: Investigation, Formal analysis, Visualization, Writing – original draft, Writing – review & editing. Jeong-A Kim: Writing – original draft, Writing – review & editing. Keun-Woo Lee: Investigation, Methodology, Data curation. Soyee Kim: Investigation, Methodology, Data curation. Kwang-Hyun Park: Writing – review & editing. Eui-Jeon Woo: Conceptualization, Supervision, Project administration, Funding acquisition. Kun-Soo Kim: Conceptualization, Supervision, Project administration, Funding acquisition.

## Data availability

The data that support the findings of this study are available from the corresponding author upon reasonable request.

## References

Abramson J et al. (2024) Accurate structure prediction of biomolecular interactions with AlphaFold 3. Nature 630:493–500 doi: 10.1038/s41586-024-07487-w

Baker-Austin C et al. (2018) Vibrio spp. infections. Nat Rev Dis Primers 4:8 doi: 10.1038/s41572-018-0005-8

DiRita VJ, Mekalanos JJ (1991) Periplasmic interaction between two membrane regulatory proteins, ToxR and ToxS, results in signal transduction and transcriptional activation. Cell 64:29–37 doi: 10.1016/0092-8674(91)90206-e

Eberhardt J, Santos-Martins D, Tillack AF, Forli S (2021) AutoDock Vina 1.2.0: New Docking Methods, Expanded Force Field, and Python Bindings. J Chem Inf Model 61:3891– 3898 doi: 10.1021/acs.jcim.1c00203

Goo SY et al. (2006) Identification of OmpU of Vibrio vulnificus as a fibronectin-binding protein and its role in bacterial pathogenesis. Infect Immun 74:5586–5594 doi: 10.1128/iai.00171-06

Gubensak N et al. (2023) Vibrio cholerae’s ToxRS bile sensing system. Elife 12 doi: 10.7554/eLife.88721

Gubensak N et al. (2021) The periplasmic domains of Vibriocholerae ToxR and ToxS are forming a strong heterodimeric complex independent on the redox state of ToxR cysteines. Mol Microbiol 115:1277–1291 doi: 10.1111/mmi.14673

Jumper J et al. (2021) Highly accurate protein structure prediction with AlphaFold. Nature 596:583–589 doi: 10.1038/s41586-021-03819-2

Kim IH et al. (2018) Cyclo-(l-Phe-l-Pro), a Quorum-Sensing Signal of Vibrio vulnificus, Induces Expression of Hydroperoxidase through a ToxR-LeuO-HU-RpoS Signaling Pathway To Confer Resistance against Oxidative Stress. Infect Immun 86 doi: 10.1128/iai.00932-17

Kim IH et al. (2013a) Transcriptomic analysis of genes modulated by cyclo(L-phenylalanine-L-proline) in Vibrio vulnificus. J Microbiol Biotechnol 23:1791–1801 doi: 10.4014/jmb.1308.08068

Kim IH, Wen Y, Son JS, Lee KH, Kim KS (2013b) The fur-iron complex modulates expression of the quorum-sensing master regulator, SmcR, to control expression of virulence factors in Vibrio vulnificus. Infect Immun 81:2888–2898 doi: 10.1128/iai.00375-13

Kim JA et al. (2015) Stationary-phase induction of vvpS expression by three transcription factors: repression by LeuO and activation by SmcR and CRP. Mol Microbiol 97:330– 346 doi: 10.1111/mmi.13028

Kovach ME et al. (1995) Four new derivatives of the broad-host-range cloning vector pBBR1MCS, carrying different antibiotic-resistance cassettes. Gene 166:175–176 doi: 10.1016/0378-1119(95)00584-1

Lee SE et al. (2000) Vibrio vulnificus has the transmembrane transcription activator ToxRS stimulating the expression of the hemolysin gene vvhA. J Bacteriol 182:3405–3415 doi: 10.1128/jb.182.12.3405-3415.2000

Martinez-Hackert E, Stock AM (1997) Structural relationships in the OmpR family of winged-helix transcription factors. J Mol Biol 269:301–312 doi: 10.1006/jmbi.1997.1065

Midgett CR, Almagro-Moreno S, Pellegrini M, Taylor RK, Skorupski K, Kull FJ (2017) Bile salts and alkaline pH reciprocally modulate the interaction between the periplasmic domains of Vibrio cholerae ToxR and ToxS. Mol Microbiol 105:258–272 doi: 10.1111/mmi.13699

Midgett CR, Swindell RA, Pellegrini M, Jon Kull F (2020) A disulfide constrains the ToxR periplasmic domain structure, altering its interactions with ToxS and bile-salts. Sci Rep 10:9002 doi: 10.1038/s41598-020-66050-5

Miller JH (1972) Experiments in Molecular Genetics. Cold Spring Harbor Laboratory

Miller VL, Taylor RK, Mekalanos JJ (1987) Cholera toxin transcriptional activator toxR is a transmembrane DNA binding protein. Cell 48:271–279 doi: 10.1016/0092-8674(87)90430-2

O’Boyle NM, Banck M, James CA, Morley C, Vandermeersch T, Hutchison GR (2011) Open Babel: An open chemical toolbox. J Cheminform 3:33 doi: 10.1186/1758-2946-3-33

Ottemann KM, Mekalanos JJ (1996) The ToxR protein of Vibrio cholerae forms homodimers and heterodimers. J Bacteriol 178:156–162 doi: 10.1128/jb.178.1.156-162.1996

Park DK et al. (2006) Cyclo(Phe-Pro) modulates the expression of ompU in Vibrio spp. J Bacteriol 188:2214–2221 doi: 10.1128/JB.188.6.2214-2221.2006

Park NY, Cho YB, Kim OB, Kim KS (2020) Cyclo(Phe-Pro) produced by Vibrio species passes through biological membranes by simple diffusion. Appl Microbiol Biotechnol 104:6791–6798 doi: 10.1007/s00253-020-10646-4

Park NY et al. (2019) Multi-Factor Regulation of the Master Modulator LeuO for the Cyclic-(Phe-Pro) Signaling Pathway in Vibrio vulnificus. Sci Rep 9:20135 doi: 10.1038/s41598-019-56855-4

Schrodinger, LLC (2015) The PyMOL Molecular Graphics System, Version 1.8. In:

Simon R, Priefer U, Pühler A (1983) A Broad Host Range Mobilization System for In Vivo Genetic Engineering: Transposon Mutagenesis in Gram Negative Bacteria. Bio/Technology 1:784–791 doi: 10.1038/nbt1183-784

Trott O, Olson AJ (2010) AutoDock Vina: improving the speed and accuracy of docking with a new scoring function, efficient optimization, and multithreading. J Comput Chem 31:455–461 doi: 10.1002/jcc.21334

