## Supplementary_Figure_1 for "Cyclic-Phe-Pro Binds the ToxRS Interface to Promote Signal Transduction in *Vibrio vulnificus*"

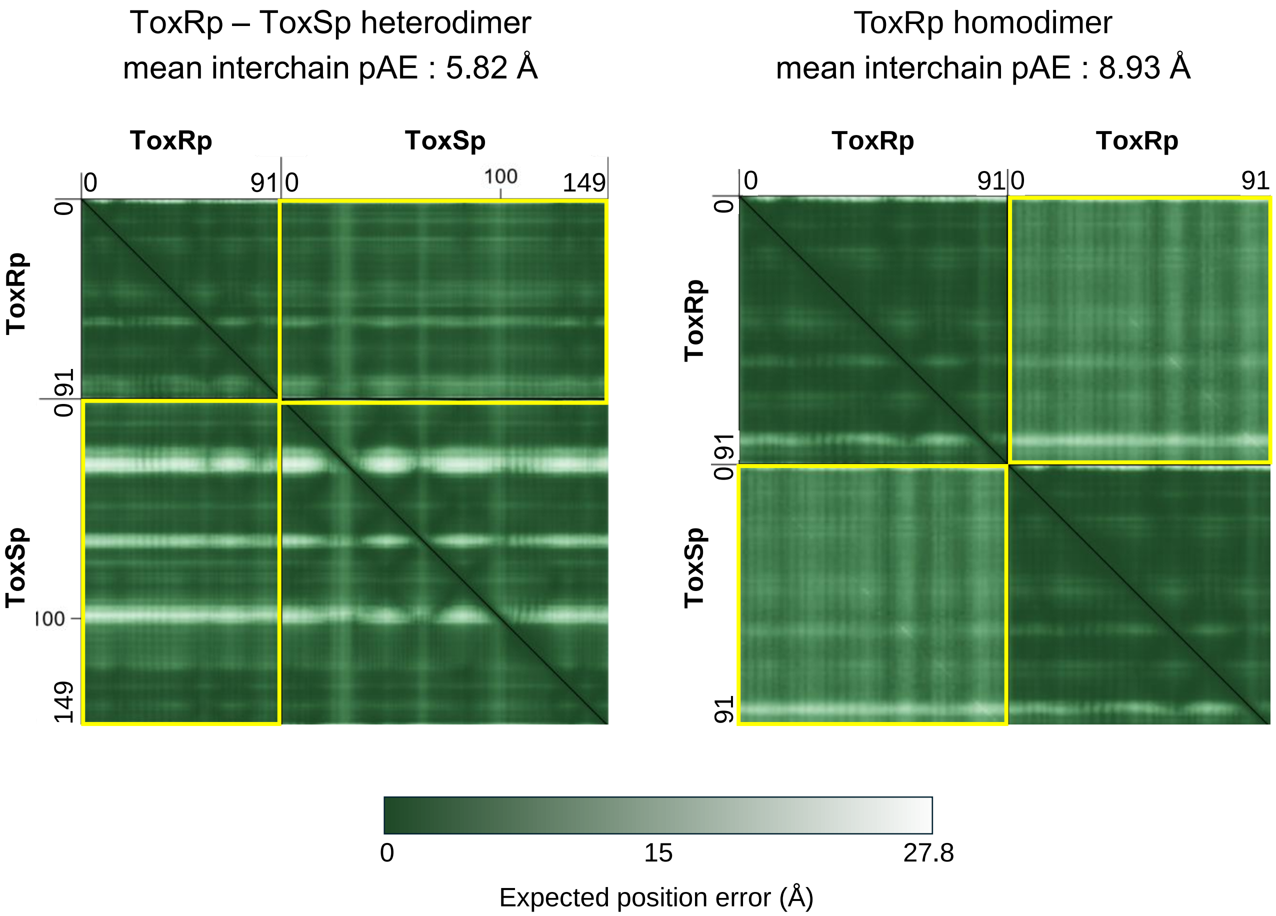


**Figure S1. Predicted aligned error (PAE) maps for the *V. vulnificus* ToxRp/Sp heterodimer and ToxRp/Rp homodimer.**
PAE matrices generated by AlphaFold3 for the ToxRp/Sp heterodimer (left) and the ToxRp/Rp homodimer (right), visualized using PAE Viewer (https://pae-viewer.uni-goettingen.de/). Each axis lists the residue positions of the two chains, separated by the vertical and horizontal lines: ToxRp (residues 1–91) and ToxSp (residues 1–149) for the heterodimer, and two ToxRp chains (residues 1–91 each) for the homodimer. The color scale (bottom) encodes the expected positional error between each residue pair, with darker shading corresponding to lower error and higher confidence. On-diagonal blocks report intrachain error, whereas the off-diagonal blocks (yellow boxes) report interchain error and reflect confidence in the predicted relative orientation of the two chains. The interchain regions of the ToxR/S map show lower PAE (mean interchain pAE 5.82 Å) than those of the ToxR/R map (8.93 Å), indicating a more confident and well-defined interface for the heterodimer than for the homodimer.
