## Supplementary_Tables for "Cyclic-Phe-Pro Binds the ToxRS Interface to Promote Signal Transduction in *Vibrio vulnificus*"

**Table S1. Strains or plasmids used in this study.**

| Strains or plasmids | Derivation / relevant characteristics | Reference or source |
|---|---|---|
| **Strains** | | |
| ***E. coli*** | | |
| DH5α | $\lambda^-$ φ80d*lacZ* Δ*M15* Δ(*lacZYA- argF*)U169 *recA1 endA1 hsdR17* ($r_K^- m_K^-$) *supE44 thi-1 gyrA relA1* | Our collection |
| S17-1 | [C600::RP4-2 (Tc::Mu)(Km::Tn7) *thi pro hsdRM*$^+$ *recA*, $Tp^r$ | (Simon et al. 1983) |
| BL21(DE3) | $F^-$ *ompT hsdSB* ($r_B^- m_B^-$) *gal dcm* (DE3) | Novagen |
| ***V. vulnificus*** | | |
| MO6-24/O | Pathogenic clinical isolate | (Reddy et al. 1992) |
| Δ*toxRS* | Derivative of MO6-24/O with a deletion in *toxRS* | This study |
| **Plasmids** | | |
| All in One™ | TA-cloning vector, *lacZ*, f1 origin, $Ap^r$ | Biofact |
| pRK415 | IncP *ori*, broad-host-range vector; *oriT* of RP4, $Tc^r$ | (Keen et al. 1988) |
| pRKΩ*lacZ* | pRK with the promoter-less *lacZ* gene for transcriptional fusion | (Park et al. 2006) |
| pRK-*leuO*::*lacZ* | pRK with the promoter-less *lacZ* gene for transcriptional fusion | (Park et al. 2019) |
| pBBR1-MCS2 | Broad range cloning vecter, $Km^r$ | (Kovach et al. 1995) |
| pBBR12::*toxRS* | pBBR1-MCS2 with the *toxRS* operon of *V. vulnificus* | This study |
| pBBR12::*toxR$_{mt}$S* | pBBR1-MCS2 with the *toxRS* operon of *V. vulnificus* with R277L and F279A mutation | This study |
| pCold™ I | Expression vector, N-terminal His-tag, $Ap^r$ | Takara |
| pColdI::*toxRp* | pCold™ I containing 193 - 291 region of *V.* | This study |

| | | |
|---|---|---|
| | *vulnificus* ToxR | |
| pColdI::*toxR*$_{mt}$*p* | pCold™ I containing 193 - 291 region of *V. vulnificus* ToxR with R277L and F279A mutation | This study |
| pColdI::*toxSp* | pCold™ I containing 20 - 173 region of *V. vulnificus toxS* | This study |

### References


Keen NT, Tamaki S, Kobayashi D, Trollinger D (1988) Improved broad-host-range plasmids for DNA cloning in gram-negative bacteria. Gene 70:191–197 doi: 10.1016/0378-1119(88)90117-5

Kovach ME et al. (1995) Four new derivatives of the broad-host-range cloning vector pBBR1MCS, carrying different antibiotic-resistance cassettes. Gene 166:175–176. doi: 10.1016/0378-1119(95)00584-1

Park DK et al. (2006) Cyclo(Phe-Pro) modulates the expression of *ompU* in *Vibrio* spp. J Bacteriol 188:2214–2221. doi: http://doi.org/10.1128/JB.188.6.2214-2221.2006

Park NY et al. (2019) Multi-Factor Regulation of the Master Modulator LeuO for the Cyclic-(Phe-Pro) Signaling Pathway in Vibrio vulnificus. Sci Rep 9:20135 doi: 10.1038/s41598-019-56855-4

Reddy GP et al. (1992) Purification and determination of the structure of capsular polysaccharide of *Vibrio vulnificus* MO6-24. J Bacteriol 174:2620–2630. doi: http://doi.org/10.1128/jb.174.8.2620-2630.1992

Simon R, Priefer U, Pühler A (1983) A broad host range mobilization system for *in vivo* genetic engineering: transposon mutagenesis in gram negative bacteria. Nature Biotechnology 1:784–791. doi: 10.1038/nbt1183-784

**Table S2. Primers used in this study**

| Name | Nucleotide sequence (5′ to 3′) |
|---|---|
| **Construction of pBBR1-MCS2::*toxRS*** | |
| toxRS_comp_F | TAG AAC TAG TGG ATC CAA AGT TCT CTA ATT GGG TG |
| toxRS_comp_R | GCT TGA TAT CGA ATT CTA AAC GAG CAT CAG TTA GAA |
| **Site-directed mutagenesis of ToxRp** | |
| toxR_SDM_F | GTGACGCTGTTAATTGCCTCAGAGC |
| toxR_SDM_R | GTTCTCGTTTGAATGTGGAAGTGTG |
| **Expression and purification of ToxRp, ToxRmt, and ToxSp** | |
| pColdI_toxRp_F | CATATGTTGCTCACCAATCCGTC |
| pColdI_toxRp_R | TCTAGATTATTTACAGATAGAGCC |
| pColdI_toxSp_F | CATATGCTTTATTGGGGTAG |
| pColdI_toxSp_R | CTGCAGTCAGTTAGAAAACAG |
| **ChIP analysis** | |
| leuO_ChIP_F | CGGAGTGGATCTCAACCTACTGAC |
| leuO_ChIP_R | ACGCATGAAAAGCTCATCATTAAA |
